# Cell and tissue type independent age-associated DNA methylation changes are not rare but common

**DOI:** 10.1101/423616

**Authors:** Tianyu Zhu, Shijie C Zheng, Dirk S. Paul, S. Horvath, Andrew E. Teschendorff

**Affiliations:** CAS Key Laboratory of Computational Biology, CAS-MPG Partner Institute for Computational Biology, Shanghai Institute of Nutrition and Health, Shanghai Institute for Biological Sciences, University of Chinese Academy of Sciences, Chinese Academy of Sciences, 320 Yue Yang Road, Shanghai 200031, China; Cardiovascular Epidemiology Unit, Department of Public Health and Primary Care, University of Cambridge, Strangeways Research Laboratory, Cambridge, CB1 8RN, UK; Department of Human Genetics, David Geffen School of Medicine, University of California Los Angeles, CA 90095, USA; Department of Biostatistics, Fielding School of Public Healthy, University of California Los Angeles, Los Angeles, CA90095, USA; UCL Cancer Institute, Paul O’Gorman Building, University College London, 72 Huntley Street, London WC1E 6BT, United Kingdom

## Abstract

Age-associated DNA methylation changes have been widely reported across many different tissue and cell types. Epigenetic ‘clocks’ that can predict chronological age with a surprisingly high degree of accuracy appear to do so independently of tissue and cell-type, suggesting that a component of epigenetic drift is cell-type independent. However, the relative amount of age-associated DNAm changes that are specific to a cell or tissue type versus the amount that occurs independently of cell or tissue type is unclear and a matter of debate, with a recent study concluding that most epigenetic drift is tissue-specific. Here, we perform a novel comprehensive statistical analysis, including matched multi cell-type and multi-tissue DNA methylation profiles from the same individuals and adjusting for cell-type heterogeneity, demonstrating that a substantial amount of epigenetic drift, possibly over 70%, is shared between significant numbers of different tissue/cell types. We further show that ELOVL2 is not unique and that many other CpG sites, some mapping to genes in the Wnt and glutamate receptor signaling pathways, are altered with age across at least 10 different cell/tissue types. We propose that while most age-associated DNAm changes are shared between cell-types that the putative functional effect is likely to be tissue-specific.

## Introduction

Age-associated DNA methylation (DNAm) changes have been reported for a long time ^1-3^. One of the first studies to indicate that age-associated DNAm changes, termed epigenetic drift, could be largely tissue specific was a study by Christensen et al ^4^ This first study however only sampled a small percentage of the DNA methylome, was largely underpowered and did not adjust for potentially confounding cell-type heterogeneity ^5,6^. Building on an observation that DNAm over specific Polycomb Repressor Complex-2 (PRC2) promoter loci correlates with age across many different tissue-types ^7^, it was demonstrated that age-associated DNAm changes can be used to build remarkably accurate predictors of chronological age, termed epigenetic clocks ^8-11^, which also appear to work independently of tissue or cell-type ^9^.

Interestingly, a recent study has suggested however that most age-associated DNAm changes are tissue-specific ^12^ Indeed, the study concluded that, with the exception of the *ELOVL2* promoter, epigenetic drift is not shared between tissues. This is a surprising conclusion given that several pan-tissue epigenetic clocks have been constructed ^9,13,14^ It led us to investigate the tissue and cell-type specific nature of epigenetic drift in more detail. In doing so, we have identified a number of critical issues with the statistical analyses performed in ^12^, which may have led to premature conclusions. First, the study performs the primary analyses using very stringent Bonferroni-corrected thresholds. While this controls for the Family-Wise-Error-Rate (FWER), the use of a Bonferroni threshold is known to suffer from a very large False Negative Rate (FNR), i.e. to a substantial reduction in power. This is particularly pertinent because their analyses generally compare age-DMPs (aDMPs) between studies and tissues, which according to our estimates were not adequately powered. Second, the authors do not report P-values of overlap, only overlapping fractions, which does not allow the statistical significance of the overlaps to be assessed. Assessing statistical significance is important because if aDMPs are not preferentially shared between tissue-types, then the reported overlaps should not deviate significantly from random. Third, the authors perform additional analyses using an arbitrary threshold on the effect size, as an alternative to statistics and P-values to select aDMPs, to argue that the “lack of overlap of aDMPs derived from different tissues” is not due to a lack of power. As we shall show here, using only a threshold on the effect size to select aDMPs is a highly problematic procedure, because of issues like heteroscedasticity, selection bias and study-specific confounding factors. Indeed, a fourth key concern is that the analysis performed in^12^ does not always fully adjust for cell-type heterogeneity, specially in complex tissues such as buccal, kidney, brain or liver. As shown recently by us, tissues like buccal, kidney or liver are highly heterogeneous, containing a substantial amount of immune-cell infiltrates ^15^, which means that adjustment for variations in the amount of infiltrating leukocytes and other stromal cells is critically important ^5,15,16^.

To address these issues, we here provide a complementary analysis to the one presented in ^12^, adjusting for cell-type heterogeneity, and using, wherever possible, matched multi cell-type or multi-tissue EWAS data from the same individuals, since such data allows for a more objective comparison between tissues and cell-types. This new analysis demonstrates that the conclusions drawn in ^12^ are too premature, and that the evidence so far points to at least 70% of epigenetic drift being shared by at least two different cell or tissue types.

## Results

### Most age-associated DMPs are shared between blood cell subtypes

First, we aimed to demonstrate that age-associated DMPs (aDMPs) derived from different cell or tissue types do exhibit a highly statistically significant overlap. We note that a highly significant overlap would be a strong cue that aDMPs shared between relevant cell or tissue-types is the norm and not the exception. In order to avoid the confounding effect of cell-type heterogeneity we first focused on three DNAm datasets profiling purified blood cell subtypes. Specifically, we analysed Illumina 450k DNAm data of 1199 purified CD14+ monocyte and 214 purified CD4+ T-cell samples from Reynolds et al ^17^, and of 100 purified CD8+ T-cells from Tserel et al ^18^. For each dataset, we derived age-associated DMPs (aDMPs), adjusting for potential technical confounders (**Methods**). We used two different significance thresholds: a false discovery rate (FDR) < 0.05, and a Bonferroni-adjusted *P*_*bon*f_ < 0.05 thresholds. The former admits 5% of false positives but ensures that the FNR is not abnormally high, whereas the Bonferroni threshold ensures in principle no false positive but at the expense of a much larger FNR. Using an FDR < 0.05 threshold, we observed a very strong and highly statistically significant overlap of aDMPs between all 3 cell-types (**Fig.1A**). For instance, almost 4000 aDMPs were found to be shared between all 3 cell-types using an FDR < 0.05 threshold (**Fig.1A**). The high statistical significance of the overlaps remained if Bonferroni thresholds were used (**Fig.1B**). Thus, even though “only” 198 aDMPs were in common between all 3 cell-types when using Bonferroni thresholds, this number was highly significant, i.e. much higher than what would be expected if aDMPs were cell-type specific.

**Figure-1:**
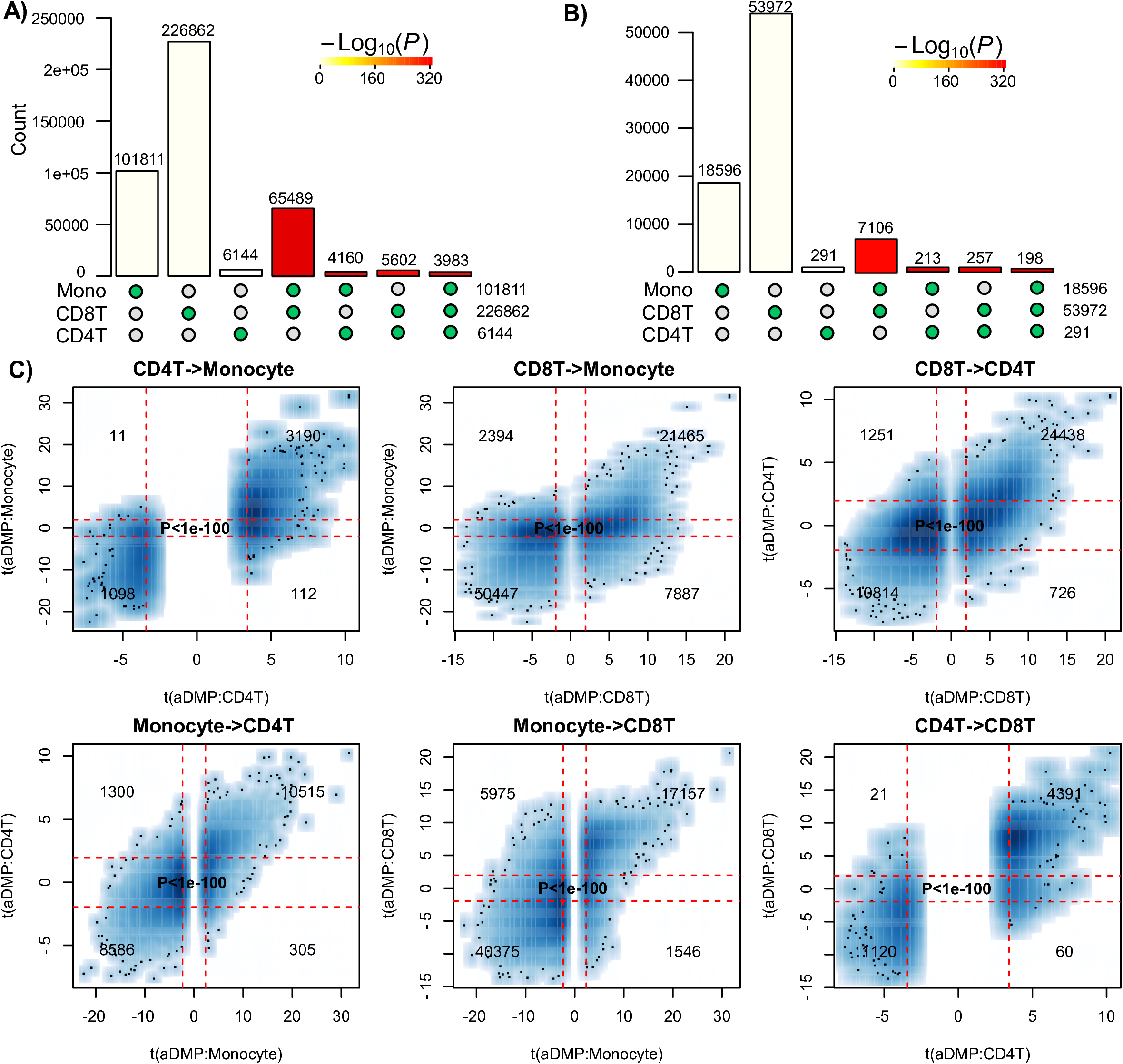
Most age-DMPs are shared between blood cell subtypes. **A)** Landscape overlap diagram for age-DMPs defined using FDR<0.05 threshold in three separate purified blood cell subtype datasets (1199 Monocytes from Reynolds et al, 214 CD4+ T-cells from Reynolds et al and 100 CD8+ T-cells from Tserel et al). Barplots indicate the number of aDMPs in each purified cell category, or the corresponding overlap between categories. For the overlapping categories, the P-value of the overlap is indicated by the color of the bar, as shown. **B)** As A), but now using a Bonferroni corrected P<0.05 threshold.

An alternative to estimating significance of overlaps, is to evaluate the consistency of the t-statistics between aDMPs, which therefore also takes into account the directionality of DNAm change. This can be done by generating scatterplots of the t-statistics of selected aDMPs in one dataset against the corresponding t-statistics in another dataset, the rationale being that if an aDMP declared in one dataset is not an aDMP in another, then its t-statistic in this other set will be distributed with a mean value of 0. Thus, if aDMPs are cell-type specific and only valid in 3 one dataset, their t-statistics in the other set profiling a different cell-type should form a data cloud centered around 0. Applying this strategy to the three purified blood cell type datasets above, revealed in each case that t-statistics were strongly skewed away from 0 and very strongly positively correlated (Fisher-test P-values <1e-100, **Fig.1C**).

In order to validate these findings, we analyzed independent data from a multi cell-type EWAS from the BLUEPRINT consortium ^19^, encompassing Illumina 450k data from CD4+ T-cells, CD14+ monocytes and CD16+ neutrophils for a total of 139 individuals. We note that the matched nature of this dataset naturally adjusts for age-range and sample size, since all 3 cell-types are available for each of the 139 individuals. We performed exactly the same analysis as before, which confirmed that overlaps were highly statistically significant, using either a FDR or Bonferroni-based threshold (**Fig.2A-B**), and importantly, that there was also a very strong correlation between the t-statistics of corresponding aDMPs (Fisher test P-values < 1e-100, **Fig.2C**), further supporting the view that a large fraction of aDMPs in one blood cell-type constitute aDMPs in another.

**Figure-2:**
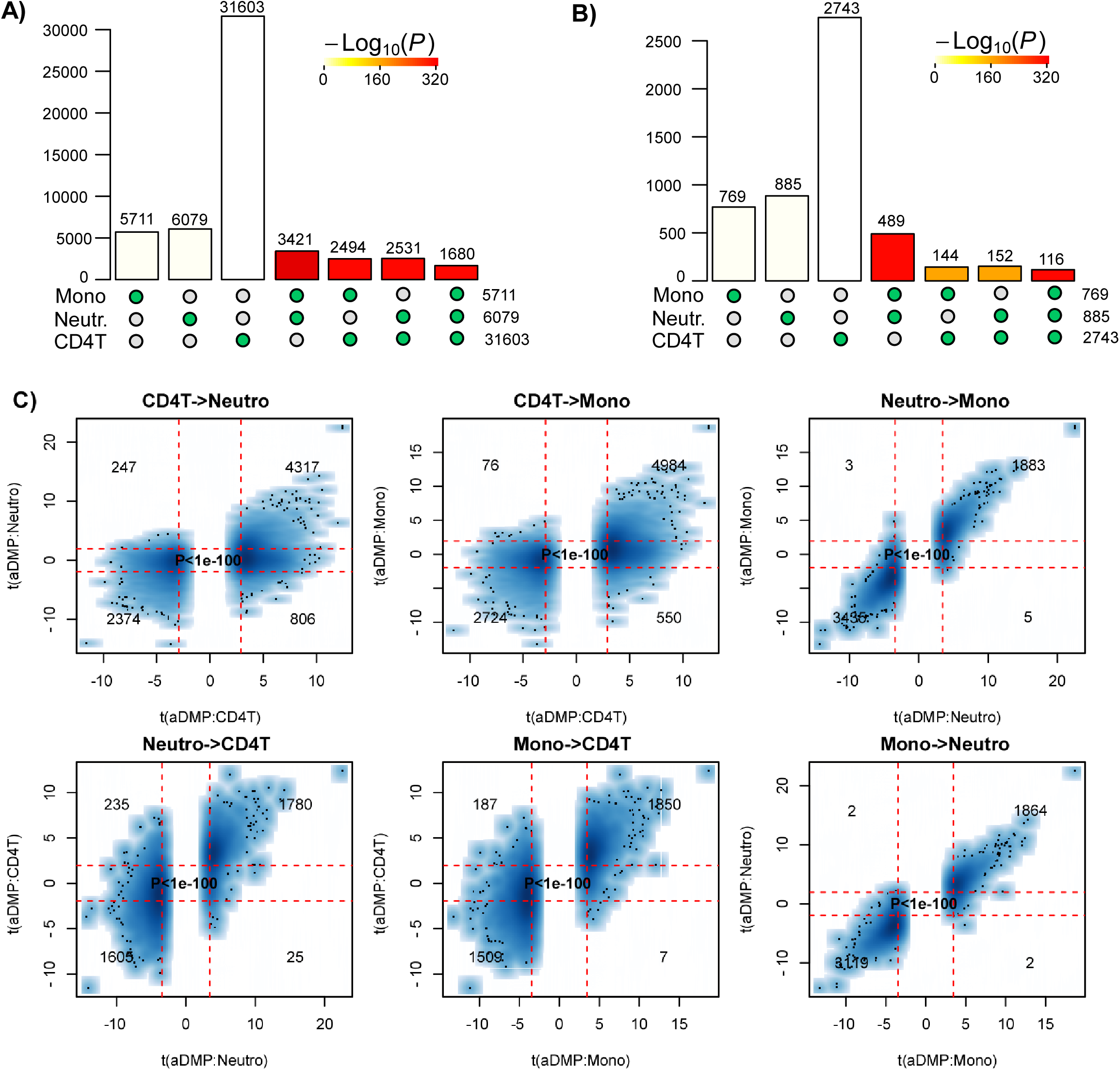
Most age-DMPs are shared between blood cell subtypes: validation in BLUEPRINT. **A)** Landscape overlap diagram for age-DMPs defined using FDR<0.05 threshold in the matched multi cell-types DNAm dataset from BLUEPRINT (139 monocyte samples, 139 matched CD4+ T-cell samples and 139 matched neutrophil samples. Barplots indicate the number of aDMPs in each purified cell category, or the corresponding overlap between categories. For the overlapping categories, the P-value of the overlap is indicated by the color of the bar, as shown. **B)** As A), but now using a Bonferroni corrected P<0.05 threshold.

### Many age-associated DMPs are shared between tissue types

Next, we decided to compare aDMPs across different tissues. These comparisons are particularly problematic if tissues derive from different datasets that profiled variable numbers of samples, with different age-ranges and subject to potentially different confounding factors, all of which could greatly impact on statistical significance estimates ^20^. Thus, ideally, a cross-tissue comparison should include multiple tissue samples from the same set of individuals, all profiled as part of the same study. Therefore, we analyzed Illumina 850k data from an EWAS profiling blood, buccal and cervical samples from a common set of 263 women (**Methods**). Because blood is a complex mixture of many immune-cell (IC) subtypes, and buccal and cervical samples are highly contaminated by immune cells ^15^, we identified aDMPs in each tissue after adjustment for batch effects and cell-type heterogeneity using EpiDISH ^21^ (in the case of blood) and HEpiDISH ^15^ (in the case of buccal and cervix). Although aDMPs identified with or without cell-type correction were highly correlated (**Fig.3A**), we observed that adjusting or not for cell-type heterogeneity did have a marked impact on the number of aDMPs, and that the number of aDMPs also varied substantially between tissue-types (**Fig.3B**). Of note, using either an FDR < 0.05 or Bonferroni adjusted P-value < 0.05 thresholds, the overlap of aDMPs between the 3 tissues was highly significant (**Fig.3C**, P<1e-100), mimicking the result obtained on blood cell subtypes. For instance, we observed a total of 2200 aDMPs in common between blood, buccal and cervix, an overlap which cannot be explained by random chance (**Fig.3C**, P<1e-100). Scatterplots of t-statistics of aDMPs between tissues further supported an extremely strong correlation, suggesting that shared aDMPs between blood, buccal and cervix is the norm and not the exception (**Fig.3D-F**).

**Figure-3:**
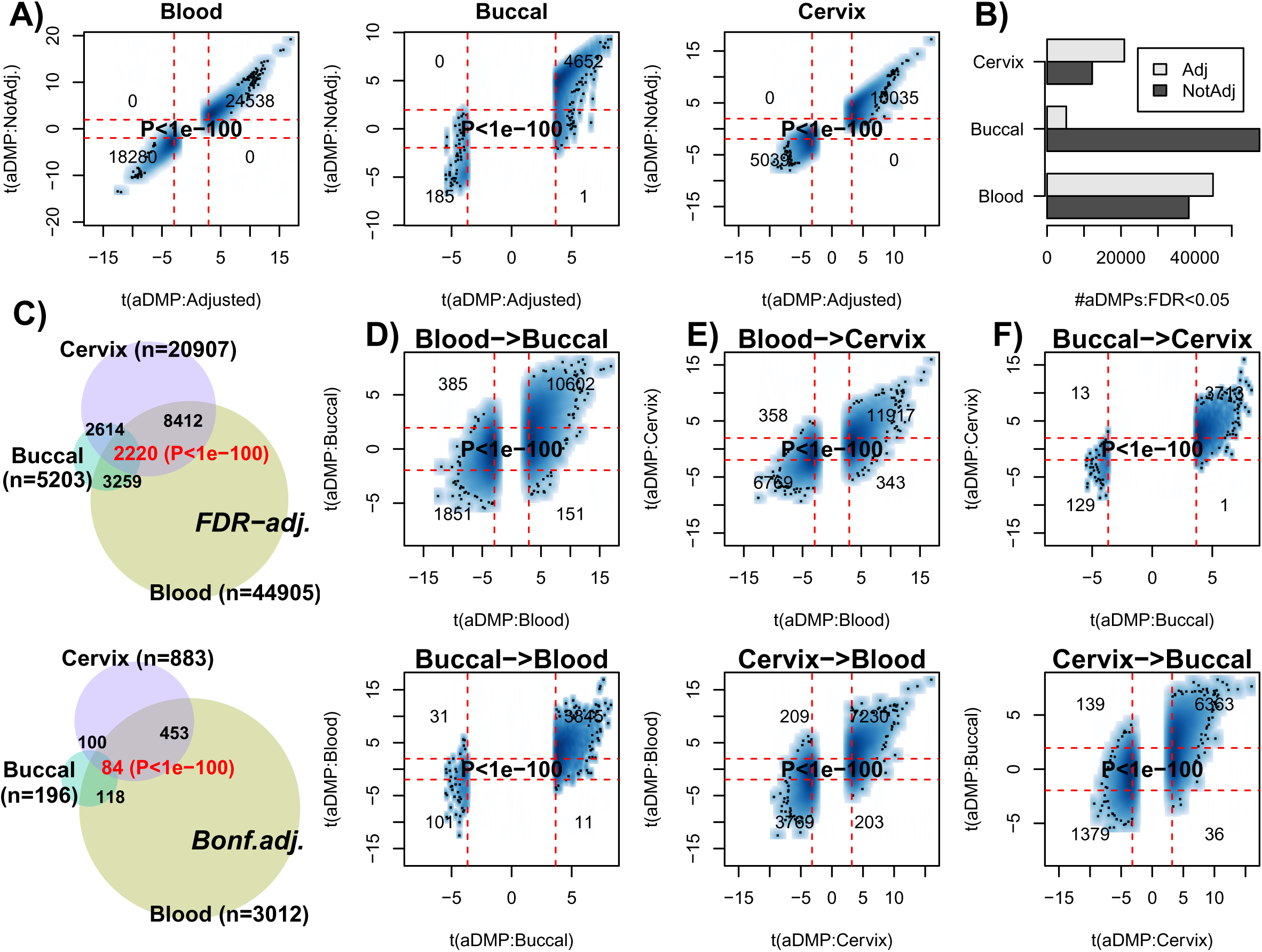
Most age-DMPs are shared between tissue-types. **A)** Smoothed scatterplots of age-DMP (aDMP) t-statistics obtained after adjustment for cell-type heterogeneity (x-axis) against the corresponding t-statistics without adjustment (y-axis), for 3 different tissue-types (Blood, Buccal and Cervix) separately. In each scatterplot, we only depict the 100 most outlier data points, we provide the number of probes in each significant quadrant and the P-value is from a one-tailed Fisher-test. The vertical red lines indicate the line of FDR<0.05, whilst the horizontal lines depict the “validation threshold” of P<0.05. **B)** Barplot of the number of aDMPs (FDR<0.05) in each tissue-type before and after adjustment for cell-type heterogeneity. **C)** Venn-diagrams representing the number of overlapping aDMPs between the 3 tissue-types using an FDR<0.05 threshold for calling aDMPs (top panel) and using a Bonferroni threshold (lower panel). P-value as estimated using a nested Hypergeometric test. **D)** Smoothed scatterplots of age-DMP (aDMP) t-statistics obtained after adjustment for cell-type heterogeneity (x-axis) in blood against their corresponding t-statistics in buccal (y-axis) for top panel, with lower panel depicting the reverse analysis, as indicated. In each scatterplot, we only depict the 100 most outlier data points, provide the number of probes in each significant quadrant and the P-value from a one-tailed Fisher-test. The vertical red lines indicate the line of FDR<0.05, whilst the horizontal lines depict the “validation threshold” of P<0.05. **E-F)** As D), but for the combinations blood-cervix and buccal-cervix, respectively.

### Studies profiling only a few hundred samples are underpowered to detect most age-DMPs

The previous analyses clearly support the view that at least thousands of aDMPs are shared between two and three tissue/cell-types. However, is this lower bound a significant underestimate on the true number of aDMPs that are shared between any two given cell or tissue types? To address this question requires careful consideration of the expected power of the studies. To estimate empirically the expected power as a function of sample size, we devised a subsampling strategy using the Reynolds monocyte dataset (**Methods**). We reasoned that this set, due to its large size (n=1199) and purified nature of the cell population, would allow us to objectively define a gold-standard set of aDMPs in monocytes. Using a stringent Bonferroni-adjusted P<0.05 threshold and using all 1199 monocyte samples, we thus defined a gold-standard set of 18596 monocyte aDMPs. We note that although this is certainly only a small subset of all true monocyte aDMPs, that it would nevertheless allow us to assess the impact of sample size on power. We next subsampled 100 samples from the 1199, and re-derived a new set of aDMPs at the same Bonferroni threshold. This subsampling strategy was performed for increasing subsampling size (from 100 to 1000, in units of 100), and a total of 10 times for each subsampling size. Sensitivity to detect the 18596 gold-standard aDMPs was computed and plotted against subsampling size, revealing that for sample sizes on the order of 100 or 200, the sensitivity was very low (**Fig.4A**). Indeed, for a subsample of 100, we estimated a mean sensitivity of only 0.00001, for a subsample of size 200 the mean sensitivity was 0.005, for 300 the value was 0.04, and at a value of 600 (i.e. about half of the full set) the sensitivity was still only 0.35. Thus, in light of this, if we were to now compare aDMPs across different tissue-types with some of the corresponding datasets in the order of 100-200 samples, as done in ^12^, then even if all aDMPs were shared between tissue-types, we would never be able to detect much overlap and we would wrongly conclude that most aDMPs are “dataset-specific”, i.e. tissue-specific in the context of the analysis presented in ^12^. For instance, using a Bonferroni threshold we observed an overlap of “only” 213 aDMPs between the 18596 gold-standard monocyte aDMPs and the 291 aDMPs derived from the 214 CD4+ T-cell samples (**Fig.1B**), and so we would be inclined to conclude that most aDMPs are cell-type specific. However, assuming that all aDMPs are shared between monocytes and CD4+ T-cells, our subsampling analysis above would suggest that the expected sensitivity to detect the 18596 gold-standard monocyte aDMPs using the 214 T-cell samples would be a value between 0.005 (n=200) and 0.04 (n=300) (**Fig.4A**), with the value closer to 0.005. Indeed, under this shared aDMP scenario, the detected overlap of 213 aDMPs represents a sensitivity fraction estimate of 213/18596 ≈ 0.1, in line with our subsampling estimate. Thus, the observed overlap of aDMPs at the given sample sizes of the two studies is not inconsistent with the great majority of aDMPs being shared between monocytes and T-cells. We note that this result is also highly congruent with the statistical significance estimates obtained previously via the Fisher-test (**Fig.1B-C**).

**Figure-4:**
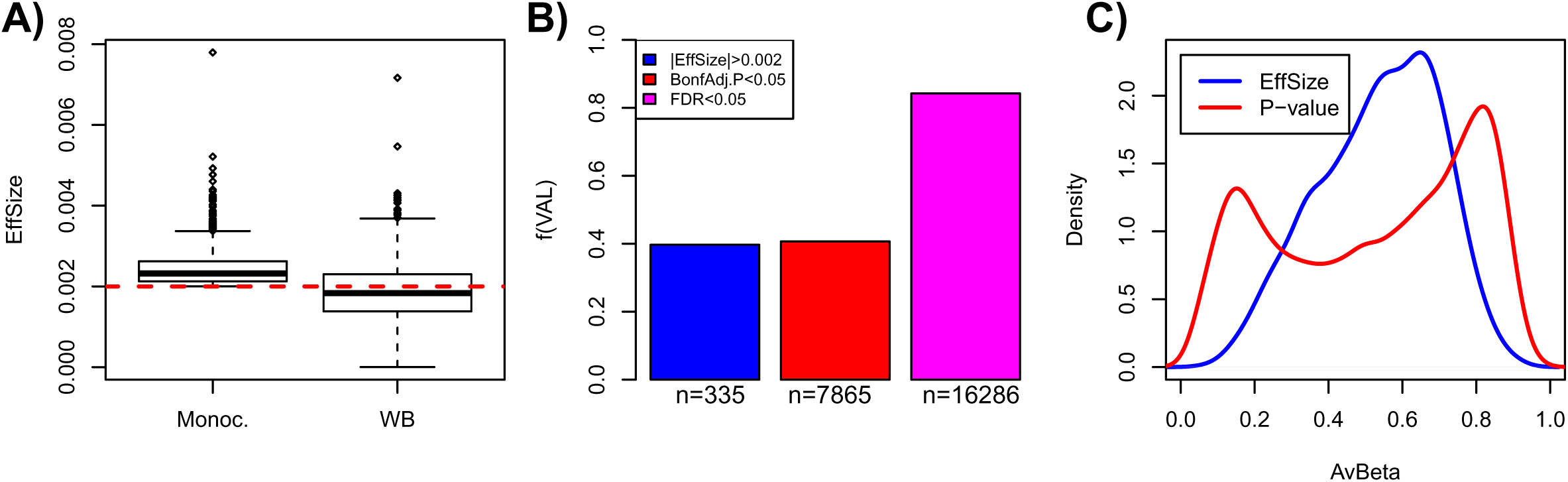
Empirical Power Analysis. **A)** Boxplot of the sensitivity (SE, y-axis) to detect gold-standard aDMPs, defined at the full purified monocyte sample size n=1199, at random subsampling sizes (x-axis), as indicated. Each boxplot displays the SE over 10 separate runs, in each run aDMPs were defined at the Bonferroni 0.05 level. **B)** As A), but now defining aDMPs in each run as those with an FDR<0.05. Because the FDR estimates are more unstable, we performed 100 runs at each subsampling size. **C)** As A), but now defining the gold-standard set of aDMPs by imposing a threshold on the effect size (2% DNAm change over 10 years), in addition to a Bonferroni adjusted P-value < 0.05. At each subsampling run, aDMPs were also defined using the same criterion and 10 runs were performed at each subsampling size. **D)** As A), but now only using the threshold on the effect size to define gold-standard aDMPs and to define aDMPs each subsampling size. A total of 100 runs at each subsampling size were performed.

We further note that using a more relaxed FDR < 0.05 threshold, sensitivity would be substantially higher: at about n=600, sensitivity is already close to 1, and for 300 samples, sensitivity is over 0.4 (**Fig.4B**). Thus, when comparing aDMPs between multiple cell or tissue-types it is even more critical to use FDR-based thresholds, since otherwise using Bonferroni-based adjustment, the expected overlap of aDMPs derived from say 4 separate studies will be zero, even if all aDMPs are common to all 4 cell/tissue-types. Our analysis suggests that many hundreds if not thousands of samples would be needed to ensure that overlaps over 3 or more studies would have the appropriate sensitivity to detect the majority of shared aDMPs (**Fig. 4A-B**). We verified that all these results are independent of whether an additional threshold on the effect size is used to select aDMPs (**Fig.4C**). Indeed, using an additional and identical threshold on the effect size to define aDMPs as used in ^12^, i.e demanding at least a 2% change in DNAm over 10 years in addition to Bonferroni significance, yielded higher sensitivities, but at sample sizes of 100 and 200, the expected sensitivity was still only 0.0002 and 0.05, respectively (**Fig.4C**).

### Using only a threshold on effect size for feature selection suffers from strong selection bias

If we were to ignore statistical significance estimates (which depend on sample size) altogether, and instead rank features by effect size using the above mentioned threshold (a 2% DNAm change over 10 years) to declare aDMPs, we can see that sensitivities increase substantially, exhibiting a much lower dependency on sample size (**Fig.4D**). For instance, at n=100, the sensitivity would be as high as 0.6 (**Fig.4D**). At first, this seems to support the argument by Slieker et al that an observed lack of overlapping aDMPs defined via an effect size threshold would imply that aDMPs are largely tissue-specific. However, this argument is incorrect, primarily for two reasons. First, selecting aDMPs based on effect size still suffers from selection bias, i.e. the fact that in the dataset where aDMPs are selected effect sizes will naturally be higher than in independent studies. This selection bias arises in real data because of numerous study-specific confounders which can significantly inflate or deflate effect sizes.

Indeed, our subsampling analysis shows that the sensitivity to detect aDMPs at a sample size of 100 to be approximately 40% lower than at the full sample size (n=1199) (**Fig.4D**), which is a substantial reduction given that the subsample derives from the same dataset. While this also demonstrates that using an effect size to rank and select aDMPs does not guarantee that the ranking and selection is independent of sample size, as claimed in ^12^, we stress that the selection bias will be even more pronounced when comparing across independent studies. Thus, the observation made by Slieker et al ^12^ that the effect sizes of aDMPs selected from one dataset appear reduced in another set profiling a different tissue could easily be the result of selection bias, and nothing to do with the nature of the different tissue being profiled.

A second major problem associated with using an effect size threshold to select and validate aDMPs is related to confounding factors such as cell-type heterogeneity, which may artificially deflate effect sizes in spite of associations with age remaining highly significant. To demonstrate this, we posited that the fraction of aDMPs derived in the monocyte set that would validate in a large whole blood set ^10^ (which contains monocytes) would be much reduced if an effect size criterion is used throughout, as compared to using a statistic and P-value. Confirming this, out of the 844 aDMPs with an effect size larger than 2% over 10 years in the Reynolds monocyte set, only about 40% passed this same threshold across the 656 whole blood samples from Hannum et al ^10^ (**Fig.5A-B**). While the fraction validating in whole blood was similar if a Bonferroni threshold is used (fraction was 41%), the fraction doubled to 84% if an FDR<0.05 threshold was used instead (**Fig.5B**). Thus, using only an effect size threshold to select and evaluate overlap of aDMPs between studies could lower sensitivities by as much as another 40% in relation to using an FDR-based threshold. Of note, using effect size thresholds in the original beta-value basis, which is highly heteroscedastic, may also strongly bias aDMPs towards those with intermediate DNAm values. Indeed, we verified that selecting aDMPs using the 2% change over 10-year threshold resulted in very few or no aDMPs with mean DNAm values close to 0 or 1 (**Fig.5C**). Since the mean DNAm level of a CpG site in a complex tissue may be strongly influenced by cell-type heterogeneity, using an effect size threshold to select or validate aDMPs could therefore easily miss a large fraction of true aDMPs.

**Figure-5:**
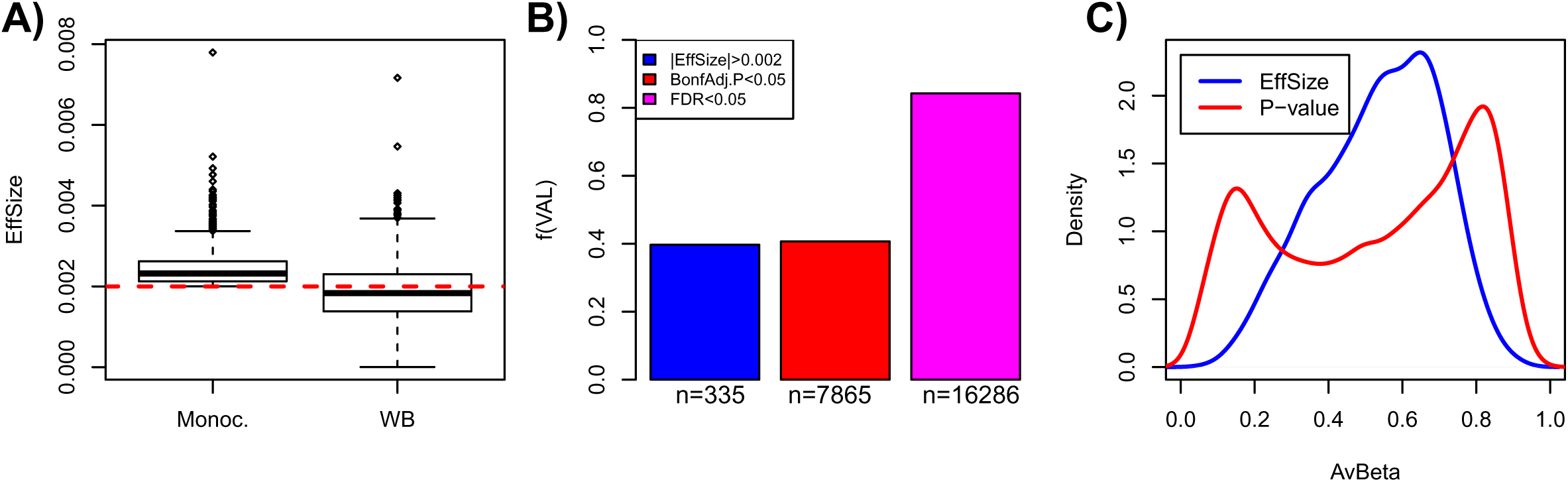
Pitfalls of using a threshold on effect-size only to select aDMPs. **A)** Boxplots comparing the effect size distribution for the 844 aDMPs defined in the Reynolds monocyte set against their effect sizes in the whole blood dataset from Hannum et al. **B)** Barplots comparing the fraction of aDMPs, defined either by the effect size threshold (blue) or P-value threshold (red & magenta, Bonferroni-adjusted), that validate in the whole blood set from Hannum et al. In Hannum et al, validated aDMPs were defined either as those passing the same effect size threshold (blue), or the same Bonferroni-threshold (red), or a more relaxed FDR<0.05 threshold (magenta). The numbers below the bars indicate the absolute number of aDMPs validating in Hannum et al. This panel demonstrates that using the same effect size threshold to define aDMPs in a dataset of complex tissue samples could miss up to 40% of true aDMPs. **C)** Comparison of the density profiles of the average DNAm for the 844 aDMPs defined by having an effect size larger (in absolute magnitude) than 0.002 (equivalent to a 2% DNAm change over 10 years) (blue line) across the 1199 monocyte samples of Reynolds et al, versus the corresponding density profile of the 19331 gold-standard aDMPs with Bonferroni adjusted P-values < 0.05.

In summary, the implicit and unproven assumption made by Slieker et al ^12^ that the effect size of an aDMP should not depend on sample size and on other factors such as cell-type heterogeneity of the tissue or other confounders, appears to be invalid and would lead to the wrong conclusion that a lack of overlapping aDMPs, all selected using effect size thresholds, is due to a lack of shared aDMPs. In fact, our analysis above clearly indicates that using only effect sizes to select aDMPs could result in a severe overall selection bias, with sensitivities reduced by as much as 80% if not more.

### FDR analysis suggests that most of the DNA methylome is altered with age

A clue as to how many aDMPs are cell/tissue specific can also be inferred from the FDR characteristics in the largest datasets. For the 1199 monocyte sample set from Reynolds et al we used the estimated FDR values (q-values) to further estimate that on average only 52419 of the 482,091 probes (i.e. 10%) are not associated with age, suggesting therefore that approximately 90% of the DNA methylome is altered with age. Because FDR estimates are notoriously sensitive to confounding factors, it is important to check the consistency of the FDR estimates in Reynolds et al against those obtained in smaller studies, ideally profiling identical or related cell-types. For the 201 monocyte samples profiled with Illumina 450k as part of BLUEPRINT, we estimated 41% of the probes to be aDMPs, whereas the fraction of aDMPs was similar, around 45%, for the 104 monocyte samples (52 twin pairs) from Paul et al ^22^ In the case of whole blood, in Hannum et al which encompassed 656 samples and therefore approximately half of the numbers in the Reynolds monocyte set, FDR values yielded an estimate of approximately 66% aDMPs. This value is very close to the one we estimated for the 689 whole blood samples profiled in Liu et al ^5^: the fraction of probes estimated there to be aDMPs was 68%. For a smaller set such as the 263 whole blood samples from FORECEE, FDR values yielded a correspondingly lower estimate of only 12% aDMPs.

Thus, assuming that the fraction of aDMPs (~90%) in the Reynolds monocyte set is inflated due to some confounder, it is unlikely to be inflated by more than 25%, since the fraction of predicted aDMPs in two large whole blood studies was consistently over 65%. This supports the view that a very high fraction (likely to be well over 65%) of the DNA methylome of blood cell subtypes is altered with age, and therefore this also means that there would be a large overlap of aDMPs between any two blood cell subtypes, consistent with our observations. Moreover, the fact that we observe equally strong overlaps of aDMPs between tissues like blood and cervix, further suggests that relatively high fractions of the DNA methylome of other cell-types (e.g. epithelial and fibroblasts) are also altered with age.

### Most of Horvath’s clock CpGs are pan-tissue aDMPs

Finally, to demonstrate that there are indeed many examples of aDMPs that are shared between tissues, we analyzed in detail the 353 CpGs that make up Horvath’s clock ^9^. Specifically, we computed their t-statistics and P-values using linear models against age, including potential covariates as confounding factors, across a total of 10 different cell/tissue types (**Methods**). Although only 3% (i.e 10 CpGs) of the 353 clock CpGs were aDMPs in all 10 tissues, approximately 78% were aDMPs across at least 3 different cell/tissue types, with only 7% appearing to be “tissue or cell-type specific” (**Fig.6**). The 10 CpGs defining aDMPs in all 10 tissue/cell types mapped to genes that included *VGF, GRIA2, FZD9, KLF14, RHBDD1, KCNC2, NHLRC1, P2RX6* and *CECR6*. Of note, *FZD9* is a transmembrane receptor for Wnt signaling proteins, whilst *GRIA2* is a glutamate neurotransmitter receptor, both of which have previously been demonstrated to be part of interactome “hotspots” of age-associated DNAm which occur independently of cell or tissue-type ^23^. Thus, this confirms that *ELOVL2* is not unique and that many of Horvath’s clock CpGs constitute aDMPs across several cell/tissue types.

**Figure-6:**
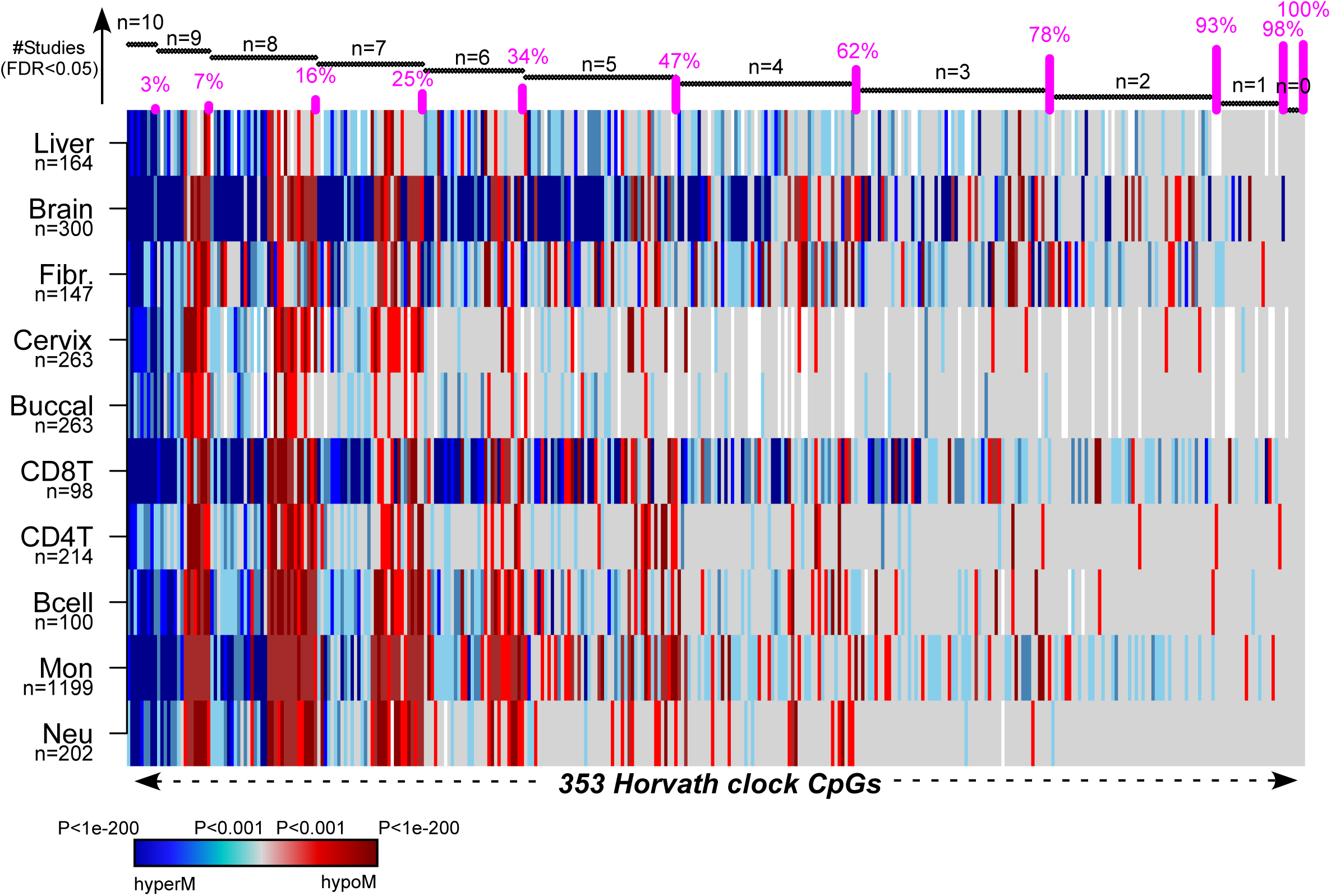
Pan-tissue analysis of Horvath’s clock CpGs. Heatmap displays the signed P-values of the 353 Horvath Clock CpGs across 10 independent cell or tissue-types, where the P-values derive from a linear model of DNAm against age plus additional confounders as covariates. Blue denotes highly significantly age-associated hypermethylation (hyperM), red denotes highly significant hypomethylation (hypoM). The 353 clock CpGs have been ranked according to the number of cell/tissue types where they are age-DMPs (using FDR<0.05), indicated by horizontal black bars at top. The cumulative proportion of the 353 CpGs that are age-DMPs in 10, 9, 8, 7, 6, 5, 4, 3, 2 or 1 studies are shown as vertical bars.

## Discussion

Here we have tried to address what appears to be an apparent paradox between a number of studies reporting pan-tissue epigenetic clocks that yield DNAm-based correlates of age independently of cell or tissue-type ^9,13,14^, and a recent study suggesting that only sites mapping to the *ELOVL2* promoter constitute cell and tissue-type independent aDMPs ^12^. While we agree with Slieker et al ^12^ that specific sites mapping to *ELOVL2* are special aDMPs in the sense that their effect sizes are particularly large across a number of different tissue-types, we strongly disagree with the other conclusion that most aDMPs are tissue-specific.

In a nutshell, their argument was based on a lack of “substantial overlap” of aDMPs derived from different tissues, either when using a very stringent Bonferroni-adjusted threshold, or when using a threshold on the effect size, which in principle is sample-size independent. In our view, their analysis is problematic, rendering their conclusions premature and potentially misleading.

First of all, to state that overlaps of aDMPs were not substantial without assessing the statistical significance of the overlaps themselves is a common pitfall not unique to their study. The expected number of overlapping aDMPs will naturally be small if a very stringent Bonferroni correction is used to define aDMPs. Even so, and as demonstrated here using several multi-tissue and multi-cell type EWAS, the overlaps using Bonferroni thresholds were highly statistically significant, a strong cue that the number of overlapping aDMPs between tissues and cell-types is not random. Indeed, using a more relaxed FDR<0.05 threshold, we have seen that there are at least thousands of aDMPs that are shared between blood cell-subtypes and also between unrelated tissues such as blood, cervix and buccal.

Second, the use of a very stringent Bonferroni threshold is specially misleading since most of the studies analysed in ^12^ were not adequately powered. Indeed, we devised an empirical subsampling analysis, which clearly demonstrated that datasets profiling only a few hundred samples or less are inadequate for assessing overlaps of aDMPs. If using Bonferroni thresholds, our power analysis suggests that many hundreds if not a thousand samples in individual studies are needed to achieve the desired high sensitivities across independent studies. We stress that these power estimates are true even for studies profiling the same cell or tissue-type, and therefore to reject the null hypothesis that most aDMPs are cell-type independent would require studies at least as large as these. That many hundreds if not thousands of samples are needed to detect large numbers of overlapping aDMPs should not be surprising: indeed, it has long been known that the ranking of features derived from large omic datasets is extremely unstable, requiring sample sizes in the order of thousands to ensure robustness of rankings under even small sample perturbations ^24^. This applies particularly to data and phenotypes characterized by small effect sizes, and would therefore apply to the case of DNAm and age. Indeed, Principal Component Analysis (PCA) on whole blood sets has consistently revealed that age-associated components of DNAm variation are generally only ranked 5^th^ or 6^th^ ^7,25^, which typically account only for a relatively small proportion (usually around 1%) of the total DNAm data variance. Thus, overlap analysis of aDMPs between pairs, or several groups, of datasets is potentially very misleading if effect sizes are small and if specific datasets are not adequately powered.

The third key problem is the use a threshold on the effect size as the sole criterion to select aDMPs, and to subsequently argue that the lack of overlap of aDMPs is not due to lack of power. While we agree that the effect size is in principle not dependent on sample-size, the lack of overlap of aDMPs defined in this way could be due to other factors. Indeed, our subsampling analysis indicated that even when using only an effect size threshold to select aDMPs, that this could still lead to reductions in sensitivity of at least 40% in studies containing only a hundred or a few hundred samples. This “selection bias”, which naturally inflates the effect sizes of the features in the studies they were derived from in comparison to other independent studies, can be further aggravated by confounders such as cell-type heterogeneity. For instance, we have demonstrated how the sensitivity to detect aDMPs defined in a pure cell-type is reduced by as much as another 40% if assessed in a cell-type mixture such as blood. Thus, overall, using only effect size thresholds to define aDMPs across tissues or cell-types could result in sensitivities to detect shared aDMPs being reduced by as much as 80% if not more. Moreover, we have shown that selected aDMPs whose effect sizes marginally miss what is an arbitrary threshold of 2% DNAm change over 10-years in independent datasets, could still be highly significantly associated with age in these same sets.

We further note that using only an effect size to select interesting features represents a “step-back” to the very old days when microarray data was first analysed, and when using thresholds on “fold-changes” in gene-expression was a common procedure. In those days, fold-changes were used because no, or very few, replicate samples were available. It was soon realized however that improved statistical inference is achieved by ranking features by a statistic. It is therefore surprising and also extremely unlikely, that an analysis based only on effect sizes can lead to critical insight not obtainable via statistics. Of note, imposing a threshold on the effect size after selecting features by statistics is a perfectly acceptable procedure.

Another related problem of using only a threshold on effect sizes to select features and which applies specifically to DNAm data, is the heteroscedasticity of beta-values ^26^. Indeed, as shown here, performing such feature selection directly on beta-values could aggravate the selection bias even further by tuning the selection of aDMPs to those exhibiting intermediate average DNAm values. This is particularly relevant in the context of assessing tissue-specificity of aDMPs, since many of the enhancer regions that are known to be highly cell-type specific would naturally exhibit such intermediate levels of DNAm. Thus, the average DNAm of such regions may be particularly variable across studies profiling different tissues, which could lead to a high FNR and reduced power.

Irrespective of all the above statistical issues, there are other strong arguments supporting the view that shared aDMPs between cell and tissue-types is the norm and not the exception. First, is the observation that in the largest studies, FDR estimates consistently indicate that most of the DNA methylome is altered with age. Using the 1199 monocyte samples from Reynolds we estimated that over 90% of the Illumina 450k probes are aDMPs. Somewhat smaller but still relatively high fractions of approximately 65% were obtained in two separate large (n>650) whole blood studies. We further note that overlaps between aDMPs from different tissues like cervix, buccal and blood were as strong as those seen between different blood cell subtypes, a strong indication that for other cell-types, say epithelial and fibroblasts, most of the DNA methylome is altered with age (as otherwise it is unlikely that we would observe such strong overlaps). Second, that most of the DNA methylome is altered with age is also highly consistent with the report of large (>1Mb) age hypomethylated blocks where sites that are normally methylated lose DNAm, and with CpG islands contained within these blocks (where sites are normally unmethylated) gaining DNAm ^27,28^. Thus, if these large-scale age-associated DNAm alterations apply to cell-types generally, and we see no good reason why they should not, shared aDMPs must be the norm, not the exception. Third, we and others ^7,13,29,30^ have observed how specific PRC2 marked sites in the genome, which are constitutively unmethylated across many fetal tissue types, consistently acquire DNAm during aging, independently of tissue or cell-type. It could well be that these specific cell-type independent aDMPs were missed in the analysis of Slieker et al due to lack of power and the use of an effect-size threshold which would bias selection against these particular sites, since these exhibit low average DNAm. Fourth, the existence of pan-tissue epigenetic clocks which can reliably predict chronological age independently of tissue or cell-type is unlikely to happen if not for a substantial number of shared aDMPs. Indeed, we estimated that at most only 7% of the 353 CpGs making up Horvath’s clock may be tissue or cell-type specific. For other aDMPs, we estimate that at the very most only 30% to 35% are cell or tissue-type specific since this is the estimated fraction of non age-DMPs in blood cell-types, and only these could be called cell-type specific aDMPs in other non-blood cell-types. Fifth, the special status of *ELOVL2* as defining the only tissue-independent aDMPs is questionable, since according to our analysis, at least another 10 CpGs mapping to unrelated genes (e.g. *FZD9, GRIA2*) constitute aDMPs across at least 10 different cell/tissue types, with almost 80% of the 353 Horvath clock CpGs defining aDMPs across at least 3 cell/tissue types.

There are of course several caveats to our analysis, which however also apply to the study of Slieker et al ^12^. First is the lack of studies profiling thousands of purified cell-types. As our empirical power analysis strongly indicates, thousands of samples would be needed to reliably determine which loci are age and not age-associated, and sample purity is important to remove cell-type composition as a potential confounder. It should be noted that even for FACS sorted cell populations, these are still only a composite and that age-associated variations in the underlying subpopulations may account for aDMPs that survive cell-type adjustment at a coarser resolution level. Another caveat is that not all tissue and cell-types analysed here derived from the same individuals, meaning that comparisons between tissues can be problematic due to differences in age-range, age distribution, gender and other study-specific factors, all of which can affect genome-wide statistical significance estimates and confound analyses. On the other hand, our analysis did include one matched multi cell-type and one matched multi tissue-type DNAm dataset, in each case encompassing 3 different cell/tissue types from the same individuals, for which age-distribution and sex were perfectly matched.

Results derived from these matched DNAm sets were in line with those obtained using unmatched sets, suggesting that the unmatched nature of some of the datasets is not a major limitation. In future however, it will be important to profile thousands of highly purified samples from a significant number of different cell-types, all from the same individuals to rigorously establish the fraction of aDMPs that are shared between cell-types.

Finally, we remark that although our analysis points towards shared aDMPs between cell-types being the norm and not the exception, that functional effects of epigenetic drift may nevertheless be tissue-specific. A concrete example is that of CpG sites mapping to the promoter of the *HAND2* gene, a target gene of the progesterone receptor. It has been indicated that gradual age-associated epigenetic silencing of *HAND2* in the endometrial stroma could inactivate the progesterone tumor suppressor pathway, sensitizing endometrial epithelial cells to oncogenic oestrogen, thus predisposing them to carcinogenic transformation ^31^. Interestingly, *HAND2* promoter sites have also been observed to undergo hypermethylation with age in blood ^28^, yet the potential functional and biological significance of this hypermethylation in blood is unclear. Thus, while specific aDMPs may be shared between tissue-types, it is only in specific tissues or cell-types that any associated functional deregulation may be of biological and clinical significance. It will be interesting for future studies to investigate whether the example of *HAND2* could serve as a more general paradigm for how shared aDMPs may exhibit functional effects in a tissue-specific manner.

## Conclusions

In summary, our novel analysis of existing datasets suggests that aDMPs shared between different cell and tissue-types is common, and not exceptional. We estimate at most 30% of aDMPs to be cell-or-tissue-type specific.

## Data Availability

All data analyzed in this manuscript are publicly available from EGA https://ega-archive.org/ accession number EGAS00001001456 and GEO (http://www.ncbi.nlm.nih.gov/geo/) under accession numbers GSE117370, GSE56581, GSE56046 and GSE59065.

## Author Contributions

Manuscript was conceived and written by AET. Statistical analyses were performed by TZ and AET. DSP prepared data from BLUEPRINT. SH contributed results derived from the fibroblast DNA methylation dataset.

## Competing Interests

The authors declare that they have no competing interests.

## Acknowledgements

This work was supported by NSFC (National Science Foundation of China) grants, grant numbers 31571359, 31771464, 31401120, by a Royal Society Newton Advanced Fellowship (NAF project number: 522438, NAF award number: 164914), by the EU-FP7 project BLUEPRINT (282510), and by European Union’s Horizon 2020 Program (H2020/2014-2020) under grant agreement number 634570.

## Methods

### DNAm datasets

#### DNAm data from purified blood cell types

We used Illumina 450k DNAm data from Reynolds et al ^17^, encompassing DNAm profiles for 1202 purified monocyte and 214 CD4+ T-cell samples. Data was downloaded from GEO (GEO accession numbers: GSE56581, GSE56046) and further processed and normalized as described by us previously ^13^. Because of confounding by gender and race, we removed 3 monocyte samples which had unique gender/race combination, leaving a total of 1199 monocyte samples. Age range for monocytes was 44 to 83. Age range for CD4+ T-cells was 45 to 79. The CD8+ T cell data was derived from Tserel et al ^18^ (GEO accession number: GSE59065), containing 100 CD8+ T cell samples, with age ranging between 22 and 84. The raw data was normalized with BMIQ ^32^. Blueprint data was derived from the European Genome-phenome Archive (EGA accession number: EGAS00001001456, BLUEPRINT study) ^19^, containing 139 CD4+ naive T-cells, 202 Neutrophil and 201 Monocyte samples with age range between 22 and 77. For the matched multi cell-type aDMP analysis we used the 139 individuals with all 3 cell-types measured. Data was normalized as described previously ^19^

#### Multi-tissue (blood, buccal and cervix) DNAm dataset

DNA methylation data encompassing whole blood, buccal swabs and cervical smears from 272 women were obtained as part of the ethically approved FORECEE study ^33^. Briefly, samples from five different European centres were sent to UCL for storage at −80C until DNA extraction. DNA extraction was performed using a Zymo spin column system. Genome-wide DNA methylation was profiled using the Infinium MethylationEPIC BeadChips (Illumina). In the case of blood and cervical samples, 500ng of genomic DNA were bisulfite converted, whereas in the case of buccal swabs, where yields were lower, 200ng were used. Pilot data had confirmed the use of 200ng to be sufficient for reliable assay-performance. BeadChips were processed by UCL Genomics using the standard recommended protocol. DNA was hybridized to BeadChips and single nucleotide extension followed by immunohistochemistry were performed using a Freedom EVO robot (Tecan). BeadChips were subsequently imaged using the iScan Microarray Scanner (Illumina). All idat files were then processed with minfi (v.1.22) using the Illumina definition of beta-value. Using the detection P-values estimated by minfi, we first computed coverage per probe (fraction of samples with detection P-value < 0.05), removing low quality probes (coverage < 0.99) and subsequently computing coverage per sample over the good-quality probes, removing low quality samples (coverage < 0.95). The small remaining number of missing values were imputed using impute.knn (with k=5) from the impute R-package ^34^. Raw data and all idat files are available from GEO under accession number GSE117370.

#### Liver DNAm dataset

We constructed a merged Illumina 450k set by combining BMIQ normalized data from two separate studies (GSE61258 & GSE48325). The merged set was defined over 417,123 probes and 164 samples.

#### Fibroblast DNAm dataset

We used the Illumina DNAm 450k dataset from ^35^, consisting of a merged set of 147 fibroblast samples.

#### Brain DNAm dataset

We downloaded the Illumina DNAm 450k set GSE74193 ^36^ from GEO. Only control non-fetal samples were used (n=300). Probes were removed if the fraction of failed samples (p>0.01) was more than 0.25, otherwise values were imputed using impute.knn function. The resulting matrix had 473536 probes left. Subsequently type-2 probe bias was normalized with BMIQ.

### Identification of age-DMPs

In each dataset we used linear models with the DNAm value as the response variable and with age of the sample as the predictor. Depending on the dataset, linear models were run with additional covariates to account for potential confounding factors. In the case of purified cell samples, covariates included those provided by the publications which included batch or ethnicity information. In the case of Reynolds et al, age-DMPs were derived by linear regression on 482127 (CD4T) and 482091 (Monocytes) probes, with gender and race as covariates (which dominated variation as determined by a PCA). In the case of the CD8+ T-cells, age-DMPs were derived by linear regression on 472484 probes, with gender and array number as covariates (which dominated variation as determined by a PCA). In the case of Blueprint data, age-DMPs were derived by linear regression on 473719 probes, with gender and batch number as covariates (which dominated variation as determined by a PCA).

In the case of complex tissues, besides adjusting for batch effects (if these were present), we also corrected for cell-type heterogeneity. Briefly, in the case of whole blood, we used our previously validated DNAm reference matrix for blood with our EpiDISH algorithm ^21^ to obtain cell-type fraction estimates for the 7 main immune cell subtypes: neutrophils, basophils, monocytes, B-cells, NK-cells, CD4+ and CD8+ T-cells. In the case of other tissues, like buccal and cervix, we used the corresponding DNAm reference matrix from our HEpiDISH algorithm ^15^ to obtain cell-type fractions for the total epithelial, total fibroblast and the 7 main immune cell subtypes. In the case of liver, we derived aDMPs using sex, body-mass index, cohort and cell-type fractions as covariates. In each case, the estimated cell-type fractions were used as covariates in the linear models.

Age-DMPs (aDMPs) were generally defined at two distinct thresholds: at a false discovery rate (FDR) threshold less than 0.05, where FDR values were estimated using the *q*-*value* procedure ^37^, and using a much more conservative Bonferroni threshold (P < 0.05/*n* with *n* the number of probes for which the linear model was run).

### Subsampling power analysis in Reynolds Monocyte set

We used the large (n=1199) purified monocyte sample set from Reynolds et al to define a gold-standard list of 19331 monocyte aDMPs using a Bonferroni threshold. We then subsampled 100, 200, 300, 400, 500, 600, 700, 800, 900, 1000 samples from the original 1199 and redefined aDMPs at each subsampling size using the same Bonferroni threshold. At each subsample size we estimated the sensitivity to detect the 18596 aDMPs from the full set. A total of 10 different runs were performed at each subsample size. We also derived aDMPs and sensitivities at each subsample size but now using an FDR<0.05 threshold. Because the FDR estimation is more unstable, we performed 100 different Monte-Carlo runs at each subsampling size.

The whole analysis above was repeated by defining aDMP by the additional requirement, that the effect size (i.e. slope) is larger than 2% over 10 years, i.e. a slope value of absolute magnitude larger than 0.002, which is the effect size threshold used in ^12^. In a final analysis, we repeated the procedure but now defining aDMPs using only the threshold on the effect size, discarding statistics and P-values.

## References

1. Ahuja, N., Li, Q., Mohan, A.L., Baylin, S.B. & Issa, J.P. Aging and DNA methylation in colorectal mucosa and cancer. Cancer Res 58, 5489–94 (1998).

2. Ahuja, N. & Issa, J.P. Aging, methylation and cancer. Histol Histopathol 15, 835–42 (2000).

3. Fraga, M.F. et al. Epigenetic differences arise during the lifetime of monozygotic twins. Proc Natl Acad Sci U S A 102, 10604–9 (2005).

4. Christensen, B.C. et al. Aging and environmental exposures alter tissue-specific DNA methylation dependent upon CpG island context. PLoS Genet 5, e1000602 (2009).

5. Liu, Y. et al. Epigenome-wide association data implicate DNA methylation as an intermediary of genetic risk in rheumatoid arthritis. Nat Biotechnol 31, 142–7 (2013).

6. Houseman, E.A. et al. DNA methylation arrays as surrogate measures of cell mixture distribution. BMC Bioinformatics 13, 86 (2012).

7. Teschendorff, A.E. et al. Age-dependent DNA methylation of genes that are suppressed in stem cells is a hallmark of cancer. Genome Res 20, 440–6 (2010).

8. Weidner, C.I. et al. Aging of blood can be tracked by DNA methylation changes at just three CpG sites. Genome Biol 15, R24 (2014).

9. Horvath, S. DNA methylation age of human tissues and cell types. Genome Biol 14, R115 (2013).

10. Hannum, G. et al. Genome-wide methylation profiles reveal quantitative views of human aging rates. Mol Cell 49, 359–67 (2013).

11. Bocklandt, S. et al. Epigenetic predictor of age. PLoS One 6, e14821 (2011).

12. Slieker, R.C., Relton, C.L., Gaunt, T.R., Slagboom, P.E. & Heijmans, B.T. Age-related DNA methylation changes are tissue-specific with ELOVL2 promoter methylation as exception. Epigenetics Chromatin 11, 25 (2018).

13. Yang, Z. et al. Correlation of an epigenetic mitotic clock with cancer risk. Genome Biol 17, 205 (2016).

14. Levine, M.E. et al. An epigenetic biomarker of aging for lifespan and healthspan. Aging (Albany NY) 10, 573–591 (2018).

15. Zheng, S.C. et al. A novel cell-type deconvolution algorithm reveals substantial contamination by immune cells in saliva, buccal and cervix. Epigenomics (2018).

16. Jaffe, A.E. & Irizarry, R.A. Accounting for cellular heterogeneity is critical in epigenome-wide association studies. Genome Biol 15, R31 (2014).

17. Reynolds, L.M. et al. Age-related variations in the methylome associated with gene expression in human monocytes and T cells. Nat Commun 5, 5366 (2014).

18. Tserel, L. et al. Age-related profiling of DNA methylation in CD8+ T cells reveals changes in immune response and transcriptional regulator genes. Sci Rep 5, 13107 (2015).

19. Chen, L. et al. Genetic Drivers of Epigenetic and Transcriptional Variation in Human Immune Cells. Cell 167, 1398–1414 e24 (2016).

20. Leek, J.T. et al. Tackling the widespread and critical impact of batch effects in high-throughput data. Nat Rev Genet 11, 733–9 (2010).

21. Teschendorff, A.E., Breeze, C.E., Zheng, S.C. & Beck, S. A comparison of reference-based algorithms for correcting cell-type heterogeneity in Epigenome-Wide Association Studies. BMC Bioinformatics 18, 105 (2017).

22. Paul, D.S. et al. Increased DNA methylation variability in type 1 diabetes across three immune effector cell types. Nat Commun 7, 13555 (2016).

23. West, J., Beck, S., Wang, X. & Teschendorff, A.E. An integrative network algorithm identifies age-associated differential methylation interactome hotspots targeting stem-cell differentiation pathways. Sci Rep 3, 1630 (2013).

24. Ein-Dor, L., Kela, I., Getz, G., Givol, D. & Domany, E. Outcome signature genes in breast cancer: is there a unique set? Bioinformatics 21, 171–8 (2005).

25. Teschendorff, A.E. et al. An epigenetic signature in peripheral blood predicts active ovarian cancer. PLoS One 4, e8274 (2009).

26. Du, P. et al. Comparison of Beta-value and M-value methods for quantifying methylation levels by microarray analysis. BMC Bioinformatics 11, 587 (2010).

27. Vandiver, A.R. et al. Age and sun exposure-related widespread genomic blocks of hypomethylation in nonmalignant skin. Genome Biol 16, 80 (2015).

28. Yuan, T. et al. An integrative multi-scale analysis of the dynamic DNA methylation landscape in aging. PLoS Genet 11, e1004996 (2015).

29. Nejman, D. et al. Molecular rules governing de novo methylation in cancer. Cancer Res 74, 1475–83 (2014).

30. Chen, Y., Breeze, C.E., Zhen, S., Beck, S. & Teschendorff, A.E. Tissue-independent and tissue-specific patterns of DNA methylation alteration in cancer. Epigenetics Chromatin 9, 10 (2016).

31. Jones, A. et al. Role of DNA Methylation and Epigenetic Silencing of HAND2 in Endometrial Cancer Development. PLoS Med 10, e1001551 (2013).

32. Teschendorff, A.E. et al. A beta-mixture quantile normalization method for correcting probe design bias in Illumina Infinium 450 k DNA methylation data. Bioinformatics 29, 189–96 (2013).

33. Widschwendter, M. et al. Epigenome-based cancer risk prediction: rationale, opportunities and challenges. Nat Rev Clin Oncol 15, 292–309 (2018).

34. Troyanskaya, O. et al. Missing value estimation methods for DNA microarrays. Bioinformatics 17, 520–5 (2001).

35. Horvath, S. et al. Epigenetic clock for skin and blood cells applied to Hutchinson Gilford Progeria Syndrome and ex vivo studies. Aging (Albany NY) 10, 1758–1775 (2018).

36. Jaffe, A.E. et al. Mapping DNA methylation across development, genotype and schizophrenia in the human frontal cortex. Nat Neurosci 19, 40–7 (2016).

37. Storey, J.D. & Tibshirani, R. Statistical significance for genomewide studies. Proc Natl Acad Sci U S A 100, 9440–5 (2003).

